# Interaction of an abiraterone with calf thymus DNA: Investigation with spectroscopic technique and modeling studies

**DOI:** 10.1101/2019.12.19.883033

**Authors:** Tanveer A. Wani, Nawaf Alsaif, Ahmed H. Bakheit, Seema Zargar, Abdurrahman A. Al-Mehizia, Azmat Ali Khan

## Abstract

Binding of toxic ligands to DNA could result in undesirable biological processes, such as carcinogenesis or mutagenesis. Binding mode of Abiraterone (ABR), a steroid drug and ctDNA(calf thymus DNA was investigated in this study using fluorescence and ultraviolet–visible spectroscopy. The probable prediction of binding and the type of interaction forces involved in the arrangement between ABR and ctDNA were explored through spectroscopic and molecular docking studies. The results indicated the binding of ABR to ctDNA in the minor groove. The binding constants were in the range of 1.35 × 10^6^ – 0.36× 10^6^ L mol^-1^ at the studied temperatures. Fluorescence and spectrophotometric data suggested static quenching between ctDNA and ABR The endothermic values of thermodynamic parameters Δ*H* = -82.8 kJ mol^−1^; Δ*S* = - 161 J mol^−1^ K^−1^ suggested that hydrogen bonding is the main force involved in binding ctDNA and ABR. In experimental studies the free binding energy at 298K was −34.9 kJ mol^−1^ with the relative binding energy ≈ −29.65 kJ mol^−1^ of docked structure. The Ksv obtained for ABR-KI was similar to that for ABR-ctDNA -KI demonstrating no protection by ctDNA against quenching effect of KI. Thus, suggesting involvement of groove binding between ABR and ctDNA. No change in the fluorescence intensity of ABR-ctDNA was observed in presence of NaCl. Thus, ruling out the involvement of electrostatic interaction. These studies could serve as new insights in understanding the mechanisms of toxicity, resistance and side effects of ABR.

## Introduction

Interaction studies of biological macromolecules with drugs are important to comprehend the mechanism involved in their interaction [1]. ABR is a pregnenolone derivative (3β)-17-(pyridin-3-yl) androsta-5,16-dien-3-ol steroid (Figure 1). ABR was developed by Institute of Cancer Research, United Kingdom. ABR is used in treatment of metastatic castration-resistant prostate cancer [2-4]. The reduced testosterone levels of <50 ng/dl have been observed after treatment with ABR in prostate cancer [5-7].

**Figure 1:**
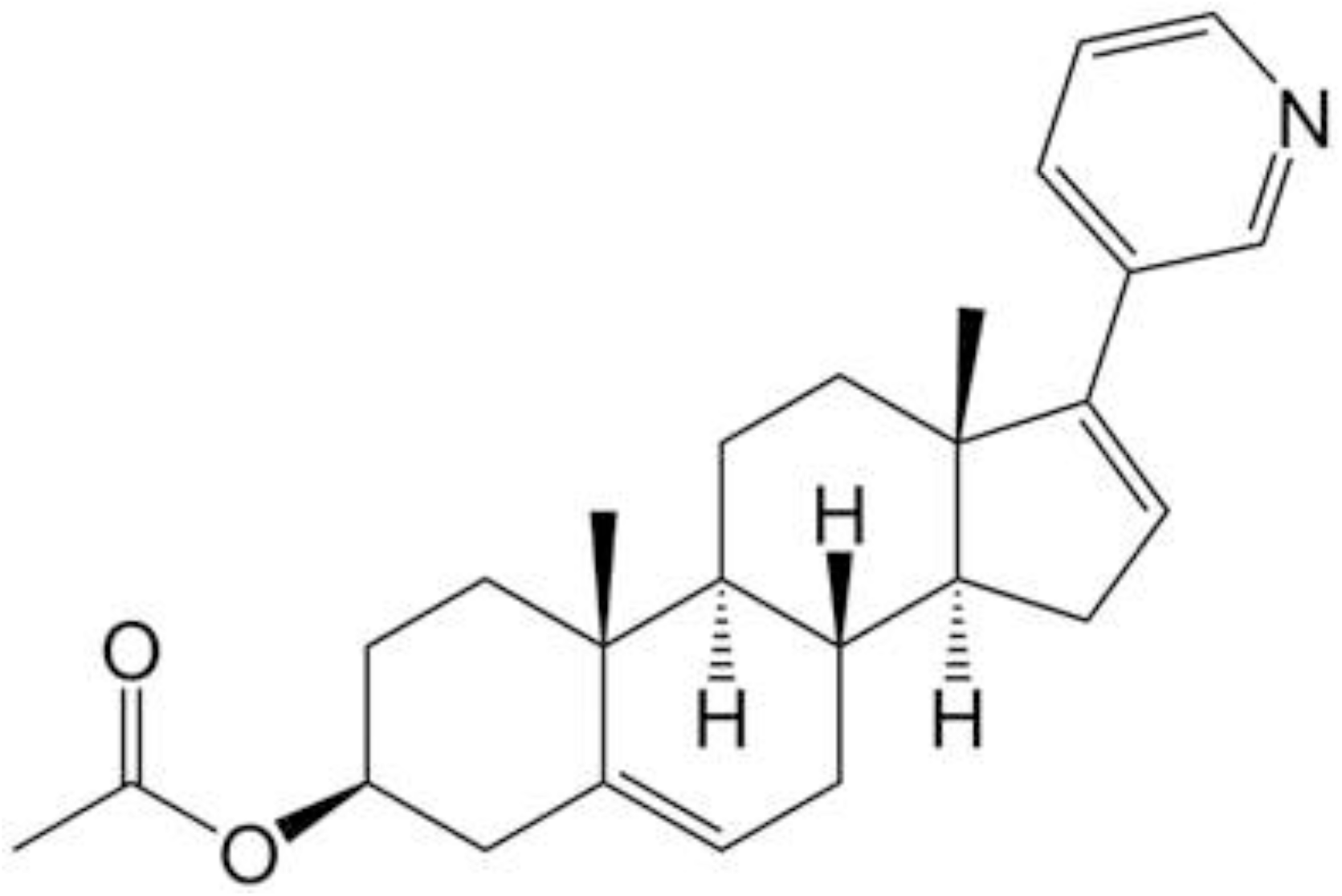
Chemical structure of ABR.

The interaction of drugs with ctDNA *in-vitro* using spectroscopic methods is currently a growing interest in research to elucidate their mechanism of action [8-11]. In general, a double stranded ctDNA could bind to the drug molecules by direct intercalation or groove binding or electrostatic interaction [8, 9]. Drugs that are capable of binding to DNA can alter the vital functions of cells by altering the expression and/or interfere in functioning of drug itself by causing drug resistance. Reactive oxygen species may also get generated because of these interactions in the DNA vicinity leading to its damage [8, 12]. Hence, the DNA-drug binding studies are valuable to understand the binding action mechanisms to DNA [13, 14]. ctDNA is preferred in these studies because of its ease in availability, structural similarities with mammalian DNA and cost effectiveness.

This study was undertaken to explore the molecular mechanisms of interaction of ABR with ctDNA and the effect of this interaction on the ctDNA itself. UV–vis absorption and fluorescence spectroscopy studies along with molecular docking analysis were performed. Further, thermodynamic studies, evaluation of quenching and binding constants, displacement studies, potassium iodide quenching studies and ionic strength studies were carried out in order to evaluate the interaction mechanism. The experimental results were corroborated with the results from molecular docking.

## Materials

ABR, CT-DNA and Ethidium bromide were purchased from Sigma Aldrich USA. Tris-HCl was procured from Himedia, India. Other reagents and chemicals used had analytical grade purity.

### Sample preparation

ABR stock solution (10mM) in ethanol and ctDNA stock (10 mg per 10 mL) in Tris-HCl 10 mM (pH 7.4) stock were prepared. The ctDNA solution was kept overnight at 4°C with intermittent shaking to make a homogenous solution. The concentration of ctDNA stock was determined with UV absorbance spectrophotometer at 260nm using molar absorption coefficient of 6600M^−1^ cm^−1^. The ctDNA purity was calculated from the A260/A280 ratio, and ratio of 1.887 indicated a protein free ctDNA. The prepared stocks were stored in dark at 4°C.

## Methods

### UV–vis spectroscopy

The UV–visible absorption spectra were acquired with Shimadzu spectrophotometer (model UV-1800, Japan). The absorption spectra (200-400 nm) for ABR and ABR bound ctDNA complex were recorded. The concentration of ABR was fixed (50 μM) with the changing concentration of ctDNA that ranged between 0–50 μM.

### Fluorescence spectroscopy-based experiments

The fluorescence spectra for ABR-ctDNA interaction were acquired with JASCO-8200 spectrofluorometer. The emission spectra of ABR (50μM) were recorded and studied in presence of ctDNA(0-50μM). The excitation wavelength (λ_ex_=265 nm) and emission wavelength (λ_em_=280-335 nm) were used to record the fluorescence spectra at (298, 303, and 310 K). Tris-HCl buffer was used to prepare the samples and the measurements were carried out in using 1.0-cm quartz cells.

### Iodide quenching studies

Iodide studies were conducted to determine the binding mode between ABR and ctDNA. Potassium iodide (KI) solutions ranging between (0-80 μm) were interacted with ABR and ABR– ctDNA separately and fluorescence intensities were recorded. Comparison of calculated Stern-Volmer constant (K_sv_) of the KI with ABR and KI with ABR-ctDNA helped in establishing the binding mode between ABR and ctDNA.

### Ionic strength studies

The fluorescence emission spectra for ABR and ABR-ctDNA were recorded in presence of sodium chloride (NaCl). Different NaCl concentrations varying from 0-70 μM were used while as, the concentrations of ABR and ctDNA were fixed.

### Conformational studies

Conformational studies circular dichroism (CD) and fourier transform infrared spectroscopic studies were conducted for ctDNA and ABR to identify any conformation changes in the protein structure on its interaction with ABR. The CD spectra for ctDNA were recorded in absence and presence of ABR. Similarly, FT IR spectra were also recorded for ctDNA in absence and presence of ABR to identify the changes in the ctDNA on its interaction with ABR.

### Molecular docking

Protein Data Bank (http://www.rcsb.org/pdb) was used to download the DNA structure (PDB ID-1BNA). For the docking analysis Molecular Operating Environment (MOE) was used and ABR structure was drawn within MOE. Default parameters MMFF94X for field force to attain energy minimization and default triangular matcher was used. London dG and GBVI/WSA dG were the scoring functions 1 and 2, respectively. The root mean square values were evaluated to find the most suitable interaction.

## Results and Discussion

### UV-visible absorption spectral studies

The absorption spectra of ABR with ctDNA are given in Figure 2. It was observed that with increasing concentration of ctDNA there was increase in the absorbance with a blue shift of 10 nm. Previous studies reported that the ligand interaction with DNA can lead to formation of complex, and the formed complex might cause red or blue shift in the absorbance of the system [9, 15]. Since an increase in the absorbance was observed, hence, intercalative binding between ctDNA and ABR is ruled out and minor groove binding between ABR and ctDNA is suggested. These findings are concomitant to previous studies of ctDNA with other drugs [16-18].

**Figure 2:**
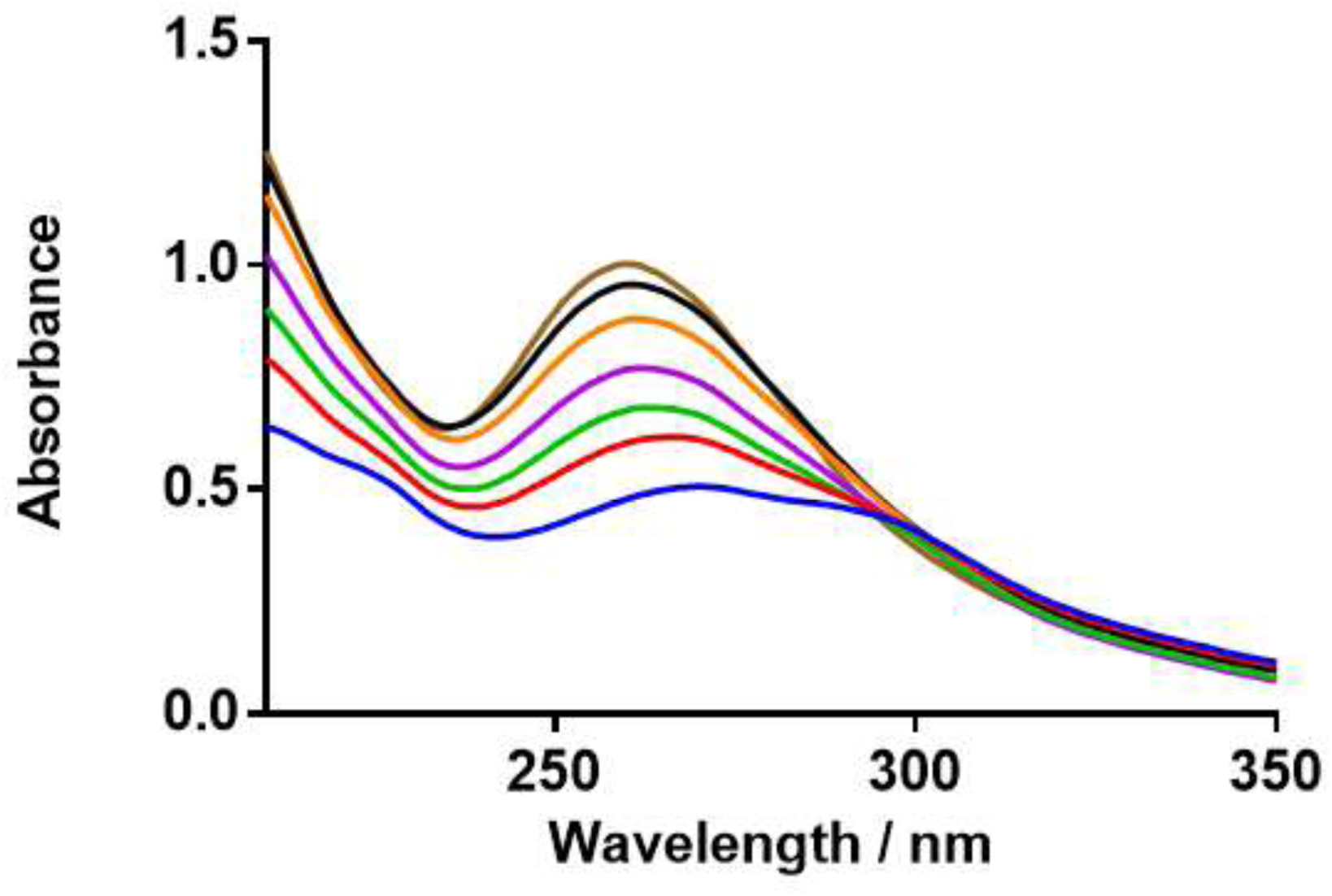
ABR UV-absorption spectra in presence and absence of ctDNA.

### Analysis on the interaction mechanism

#### Fluorescence spectrum

The mechanism of interaction between ligands and bio-macromolecules can be understood using steady state fluorescence spectroscopy. The fluorescence spectroscopy helps in computing the binding constant and provides information regarding the type of forces involved in the interaction. ABR demonstrated fluorescence emission at (λ_em_=339 nm with excitation wavelength at (λ_ex_=265 nm, whereas, ctDNA didn’t exhibit fluorescence by its own. A decrease in the fluorescence intensity of ABR was observed with addition of ctDNA to ABR. The fluorescence intensity decreased further on gradual increase in the ctDNA concentration (0-50 μM) with a fixed concentration of ABR (50 μM). The ABR and ctDNA fluorescence spectra at 298 K are given in Figure 3. Thus and interaction between ctDNA and ABR was suggested. The inner filter effects were eliminated and the fluorescence intensity was corrected [19].

**Figure 3:**
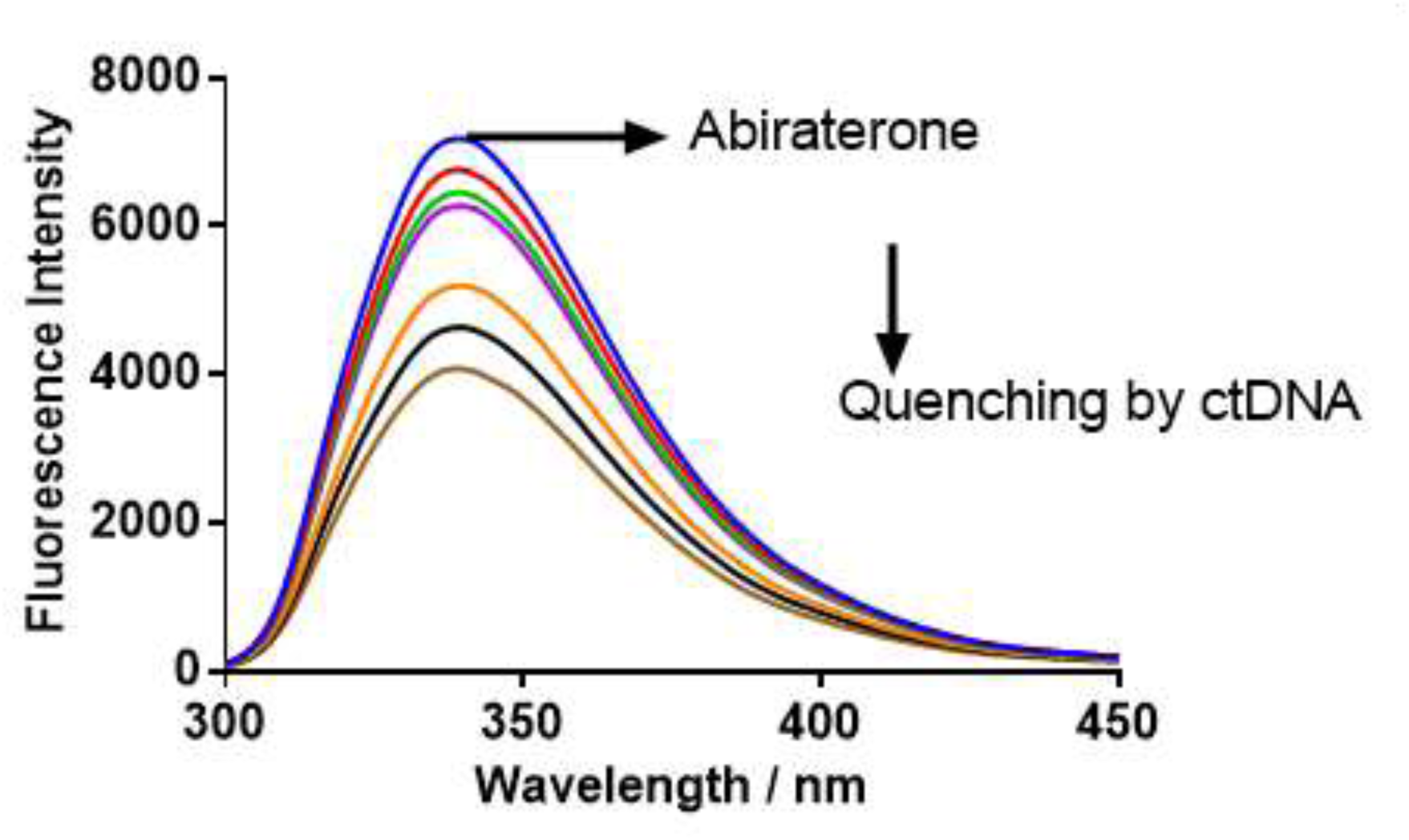
ABR fluorescence spectra in absence and presence of different concentrations of ctDNA at 298 K.

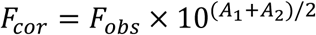

Where, F_cor_ is corrected and F_obs_ is observed fluorescence intensity; A_1_ and A_2_ are the respective absorbance at excitation and emission wavelengths.

### Quenching constant and quenching mechanism

The quenching mechanisms involved in the biomolecules and ligands can either be static or dynamic quenching and are distinguished from one another based on their behavior at different temperatures. Diffusion is the basis of dynamic quenching, higher diffusion and strong collision quenching occurs at higher temperature. The fluorophore interacts with the quencher and forms a non-fluorescent complex amongst themselves in case of static quenching. The complex formed is relatively unstable as the temperature increases in case of static quenching [20]. The quenching of fluorescence was evaluated by the Stern-Volmer equation [21] as follows:

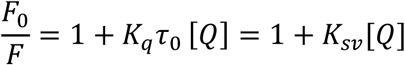

where F_0_ and F represent fluorescence intensity of fluorophore and with quencher, respectively; k_q_ is bimolecular quenching constant; τ_0_ is the lifetime of the fluorophore without quencher; [Q] is the concentration of the quencher; and K_SV_ is the Stern–Volmer quenching constant [20]. The Ksv at three different temperatures of (298, 303 and 310 K) were calculated and are presented in Figure 4A, with the *Ksv* values given in Table 1. An increase in the *Ksv* values was observed as the temperature increased. However an increase the absorbance spectrum of ABR and ctDNA was observed upon increasing the concentration of ctDNA suggests complex formation between the two.. Further, the quenching constant values obtained (Table 1) were higher than the maximum scattering collision quenching constant 2.0 × 10^10^ L mol^-1^ s^-1^. [22-25]. Thus, it was inferred that static quenching occurred in ABR –ctDNA complex.

**Table 1:** The Stern-Volmer quenching constants K_SV_ and the quenching rate constants for ABR - ctDNAinteraction

| **T(K)** | **R** | **$K_{sv}$ (L mol<sup>-1</sup>)** | **$K_q$ (L mol<sup>-1</sup>s<sup>-1</sup>)** |
| --- | --- | --- | --- |
| 298 | 0.9939 | $1.79 \times 10^4$ | $1.79 \times 10^{12}$ |
| 303 | 0.9948 | $2.00 \times 10^4$ | $2.00 \times 10^{12}$ |
| 310 | 0.9936 | $2.04 \times 10^4$ | $2.04 \times 10^{12}$ |

**Figure 4:**
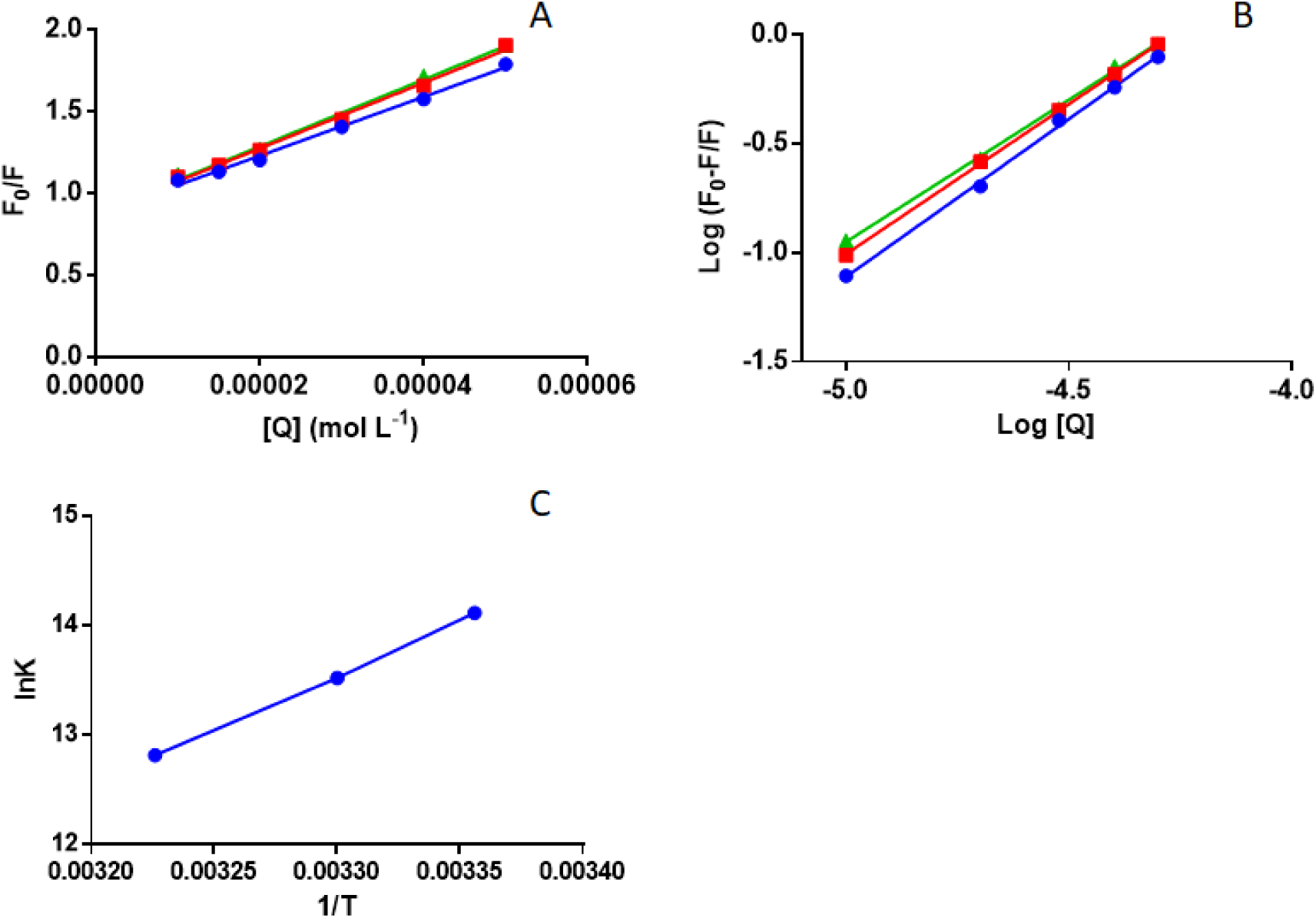
[A]: ABR Stern–Volmer plot for its quenching by ctDNA (298/303/310) K; [B]: The plot of log [(F_0_-F)/F] versus log[Q] for quenching process of ABR with ctDNA at 298/303/310 K; [C]: Van’t Hoff plots for binding interaction of ABR and ctDNA

### Binding constant and binding sites

The binding constant “K_b_” was obtained from double logarithmic regression curve. It also provided information regarding the number of binding sites “n” [25-28]. From the linear regression curve of log (F_0_−F)/F *vs* log [Q] (Figure 4B) the “Kb” and “n” values were determined [13, 22, 23]:

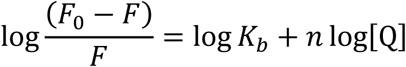

The K_b_ and n for the ABR-ctDNA systems are given in Table 2. The value (n≈1) at all the studied temperatures infer that ABR and ctDNA interacted in a molar ratio of 1:1. A decreased binding constants at high temperatures suggested ABR’s dissociation from ABR-ctDNA complex. In general terms a binding constant values in the range 10^6^ to 10^8^ M^−1^ suggested a strong binding affinity [29, 30]. The binding constant values between ABR and ctDNA(Table 2) indicate a relatively strong binding interaction.

**Table 2:** Binding parameters and thermodynamic parameters of ABR-ctDNA.

| **T(K)** | **R** | **$K_b$**<br>**( L mol<sup>-1</sup>)** | **n** | **$\Delta G$**<br>**(kJ mol<sup>-1</sup>)** | **$\Delta H$**<br>**(kJ mol<sup>-1</sup>)** | **$\Delta S$**<br>**(J mol<sup>-1</sup>K<sup>-1</sup>)** |
| --- | --- | --- | --- | --- | --- | --- |
| 298 | 0.9976 | $1.35 \times 10^6$ | 1.44 | -34.93 | -82.8 | -161 |
| 303 | 0.9983 | $0.74 \times 10^6$ | 1.37 | -34.13 | | |
| 310 | 0.9986 | $0.36 \times 10^6$ | 1.30 | -33.00 | | |

### Binding mode and interaction forces

The non-covalent interactions that could be involved in the binding of ligands and bio-macromolecules are hydrogen bonds, van der Waals forces, and hydrophobic and electrostatic interaction [31]. The types of non–covalent interactions involved can be identified with the thermodynamic parameters such as enthalpy change (ΔH°) and entropy change (ΔS°). Hence to evaluate binding mode and interactions the thermodynamic parameters were calculated by Van’t Hoff equation:

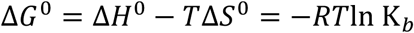

where R is the gas constant, T is the experimental temperature, and ΔG° is free energy change. A plot of *ln* K_b_ against 1/T was explored. The intercept and the slope of the plot are used to calculate ΔH° and ΔS° (Figure 4C) and the values are given in Table 2. The results suggested that hydrogen bonds contribute in the interaction between ABR and ctDNA. [8, 22]. Negative ΔG° indicated spontaneous interaction between ctDNA and ABR. The results are in concurrence with docking studies since the free binding energy at 298K was -34.9 kJ mol^−1^ and the predicted energy from molecular docking was ≈ −29.65 kJ mol^−1^. The difference in the theoretical and the experimental results can be attributed to variation in the solution environment for experimental analysis and space model for docking analysis [32].

### Ethidium Bromide (EB) competitive experiment

Ethidium bromide (EB, 3,8-diamino-5-ethyl-6-phenyl-phenanthridinium bromide) is an intercalating agent in the DNA binding [10, 33] and therefore, was used to investigate and ascertain if the intercalative mode of binding was involved in the interaction. On interaction with ctDNA, EB gives an intense emission whereas, no noticeable emission is observed with EB alone. The emission spectra for EB –ctDNA were recoded both with and without ABR at 500-700 nm (λex = 524 nm nm). However, no noticeable change was observed in the emission spectrum of EB-ctDNA on addition of ABR (Figure 5). EB was not displaced from its site of binding on ctDNA by ABR as no change in the emission of EB-ctDNA was observed suggesting non-intercalative binding of ABR and ctDNA.

**Figure 5:**
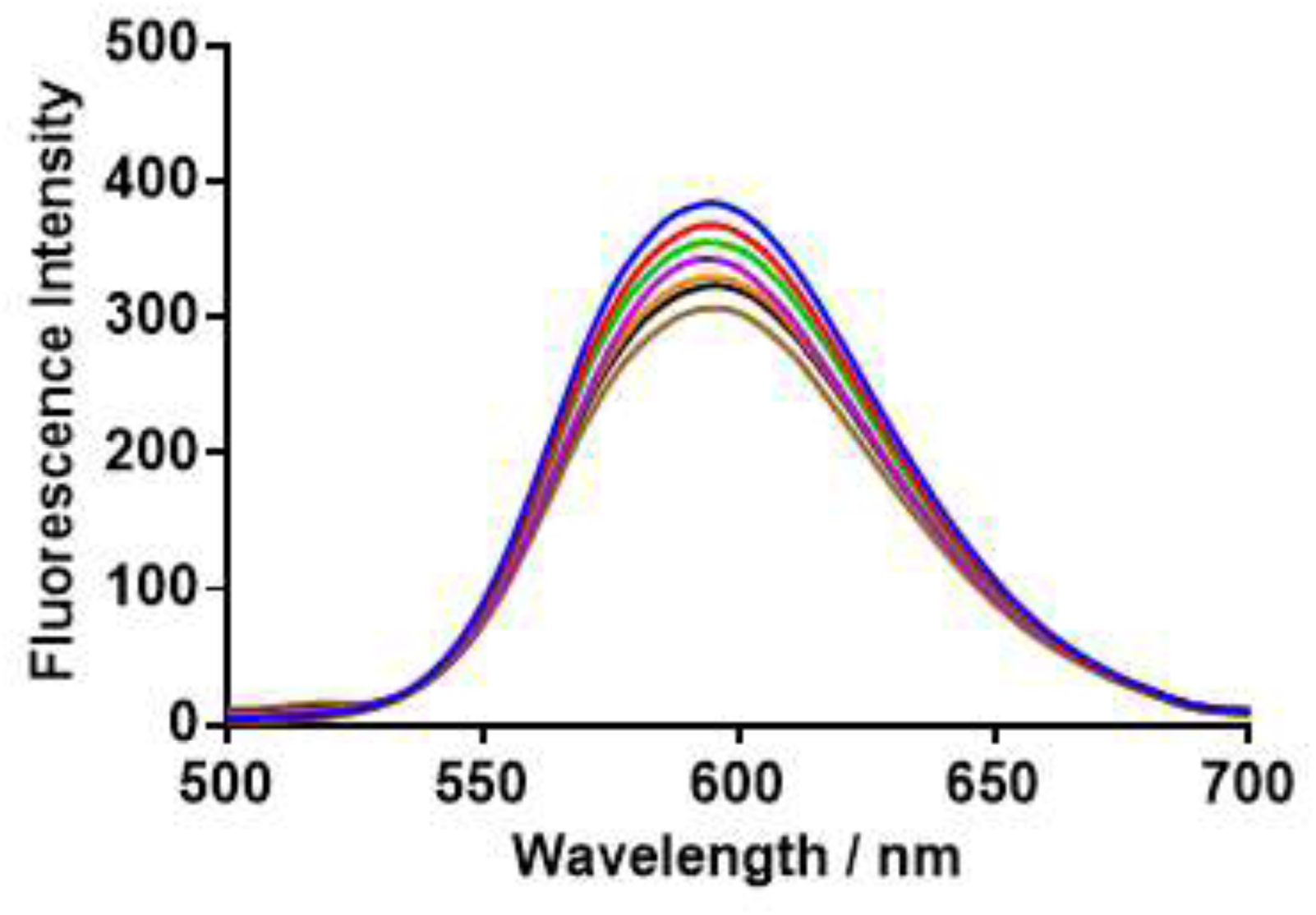
Fluorescence spectra of ctDNA and EB (intercalator) complex in presence of varying concentrations of ABR (0-50 μM).

### Iodide quenching studies

Binding interaction between ctDNA and ABR were further elucidated with iodide quenching studies. Potassium iodide (KI) was used as quencher for ABR and ABR ctDNA. In case of intercalative interaction between ligand and ctDNA the ligand is protected by the double stranded ctDNA from the quenching effect of ionic quencher whereas, in case of groove binding between ligand and ctDNA the quenching should occur readily as the ligand is not protected from the effect of anionic quencher[34]. The quenching constants (Ksv) for ABR and ABR-ctDNA with KI were calculated using Stern-Volmer equation Figure 6. The Ksv obtained for ABR-KI was similar to that for ABR-ctDNA -KI demonstrating no protection by ctDNA against quenching effect of KI. Thus, suggesting involvement of groove binding between ABR and ctDNA[13].

**Figure 6:**
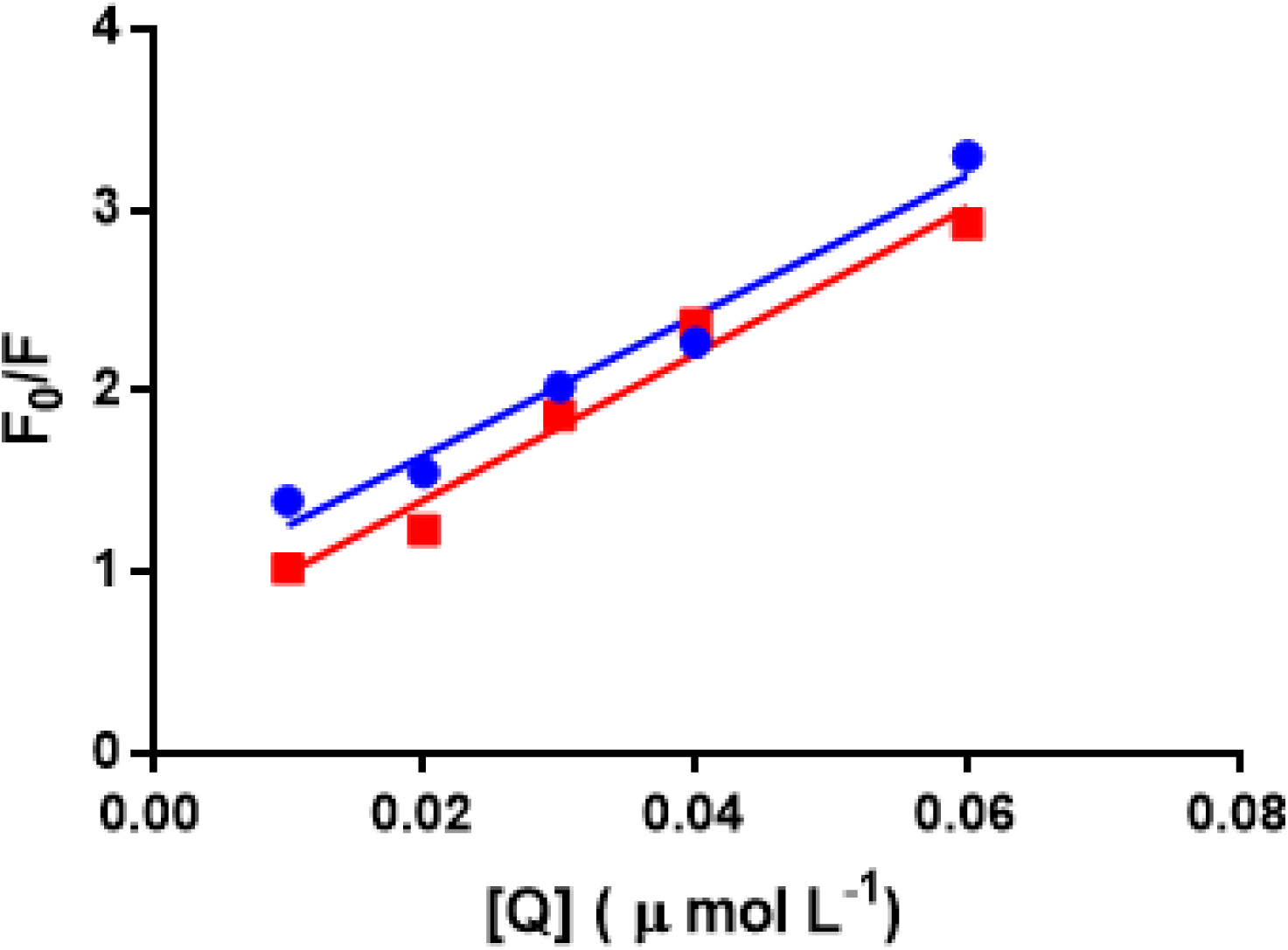
Stern-Volmer plot for fluorescence quenching of ABR by KI in absence and presence of ctDNA

### Effect of Ionic Strength

The ionic strength studies were used to rule out the electrostatic binding mode between ABR and ctDNA. Sodium chloride (NaCl) is generally used for this purpose as it’s addition has minimal effect on the fluorescence intensity of the small drug molecule. The negative charge of DNA phosphate backbone is neutralized by the sodium (Na^+^), thus, decreasing the electrostatic repulsion amid weakening of surface-binding interactions. Also, the addition of Na^+^ leads to decreased electrostatic attraction amid the DNA surface and the small drug molecule. In electrostatic binding, the binding occurs outside the groove and addition of NaCl results in reduced quenching of FI [12]. In the current study no change in the fluorescence intensity of ABR-ctDNA was observed in presence of NaCl. Thus, ruling out the involvement of electrostatic binding mechanism between ctDNA and ABR.

### Circular dichroism (CD) spectra studies

Circular dichroism studies are helpful in identifying conformational changes in the protein on its interaction with ligand. Two bands can be observed in the UV region of CD spectrum of B-DNA. One of the bands due to right handed helicity appear at 245 nm (negative region) whereas the second band from base stalking appear in the positive region at 275 nm [35]. The CD spectra (Figure 7) were recorded for ctDNA (50 μM) and ABR (0, 20, 40 μM). In case of groove binding or electrostatic interaction between DNA and ligand, no alteration in the base stalking and helicity occurs. However, in intercalative interaction there is change in the base stalking and helicity of the DNA, thus altering the conformation of DNA [35].

**Figure 7:**
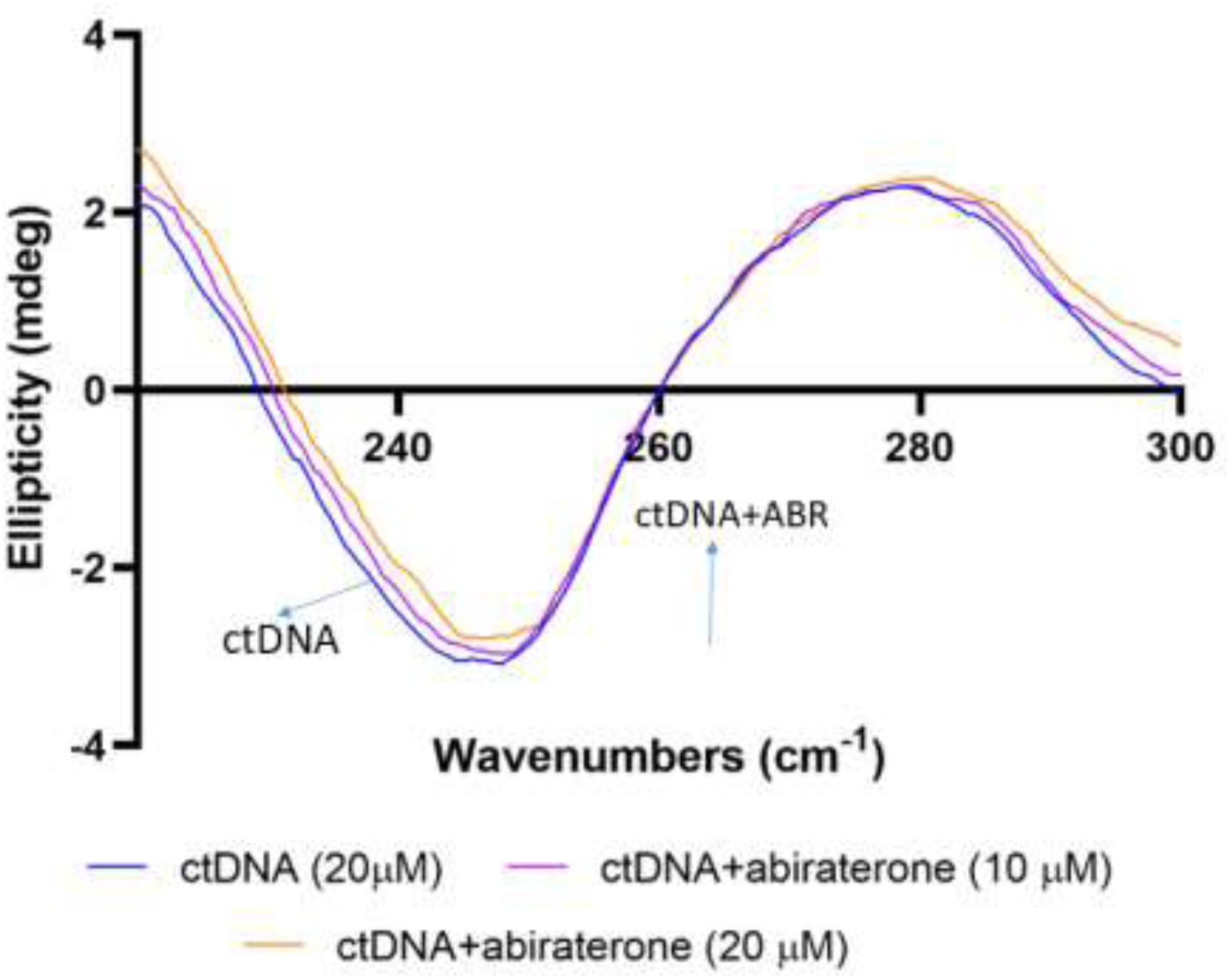
Circular dichroism spectra of ctDNA in presence and absence of ABR

The CD spectrum of ctDNA didn’t showed any significant changes in the presence of absence of ABR suggesting grove binding interaction between ctDNA and ABR. The minor changes in the bands upon ABR addition to ctDNA is attributed to nucleic acid structure transition to compact φ structure from an extended conformation. [36]

### FT-IR analysis

The FT IR is used for the characterization and structural changes in the protein on its interaction with ligand. The FT-IR analysis for ctDNA was carried out in the spectral region 1000-1800 cm^-1^ (Figure 8). The spectra were recorded for ctDNA and ctDNA-ABR. The DNA nitrogen bases of guanine (G), thymine (T), adenine (A) and cytosine (C) correspond to the vibration bands 1733, 1684, 1631 and 1520 cm^-1^, respectively.

**Figure 8:**
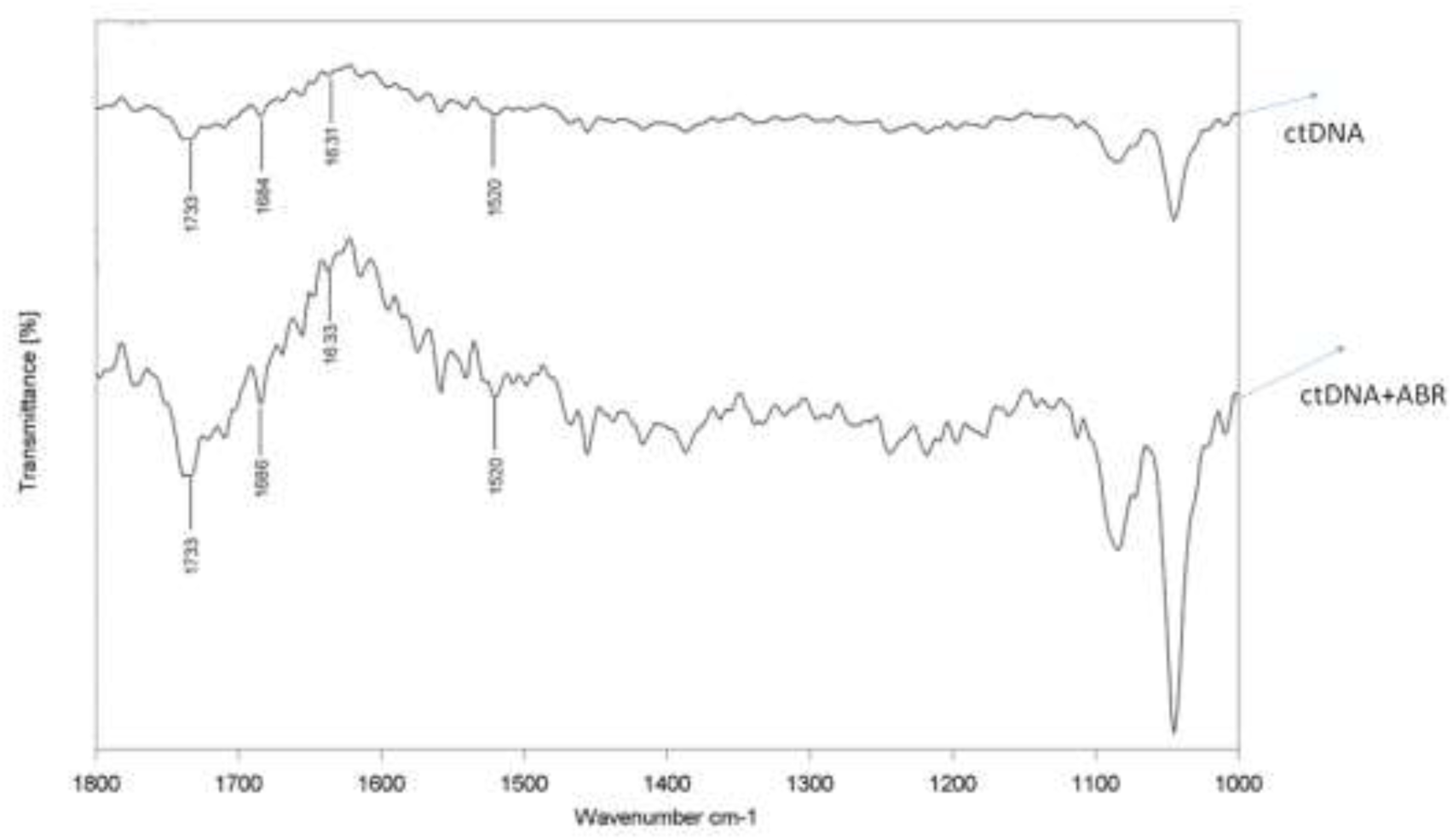
FT IR spectra of ctDNA in presence and absence of ABR

The ligand protein interaction is suggested on the basis of shift in the peaks of the FT IR spectra. The T band shifted from 1684 to 1686 cm^-1^ and the A band shifted from 1633 to 1631 cm^-1^ The changes in the position of the peak of FT-IR spectra is an indicator of the ligand –DNA interaction. In, the FT-IR spectra major shifts were seen in the T band from 1663 to 1665 cm^-1^ and for A band from 1632 to 1626 cm^-1^ were observed due to the interaction. An increase in the intensity of all nitrogen bases was observed on interaction of ctDNA and ABR. The shift in the thymine and adenine bands in ctDNA -ABR complex suggested interaction of ABR with T and A bases. Although, there was no shift in the G and C bases however, interaction with these two cannot be ruled out. The results for the FT IR analysis were in accordance with the molecular docking studies.

### Molecular modeling

Molecular docking serves as an instrumental tool in studying ligand and macromolecular interactions [13, 37]. Molecular docking using MOE for ABR and ctDNA was executed to attain an understanding in terms of the binding site and binding location for ABR in the ctDNA. The model which showed the lowest docking energy was taken for the docking. The docking results revealed that the ABR interacts with the ctDNA via minor groove binding. (Figure 9), also, the interaction was non-covalent in nature. These results are in agreement to the values obtained experimentally. The binding energy for ABR and DNA was found to be -7.088 kcal mol^-1^ with the docking analysis and experimentally it was found to be -8.34 kcal mol^-1^. Further, a hydrogen bond was also found between 3-N of ABR and 2-N of DG22 of chain B of DNA.

**Figure 9:**
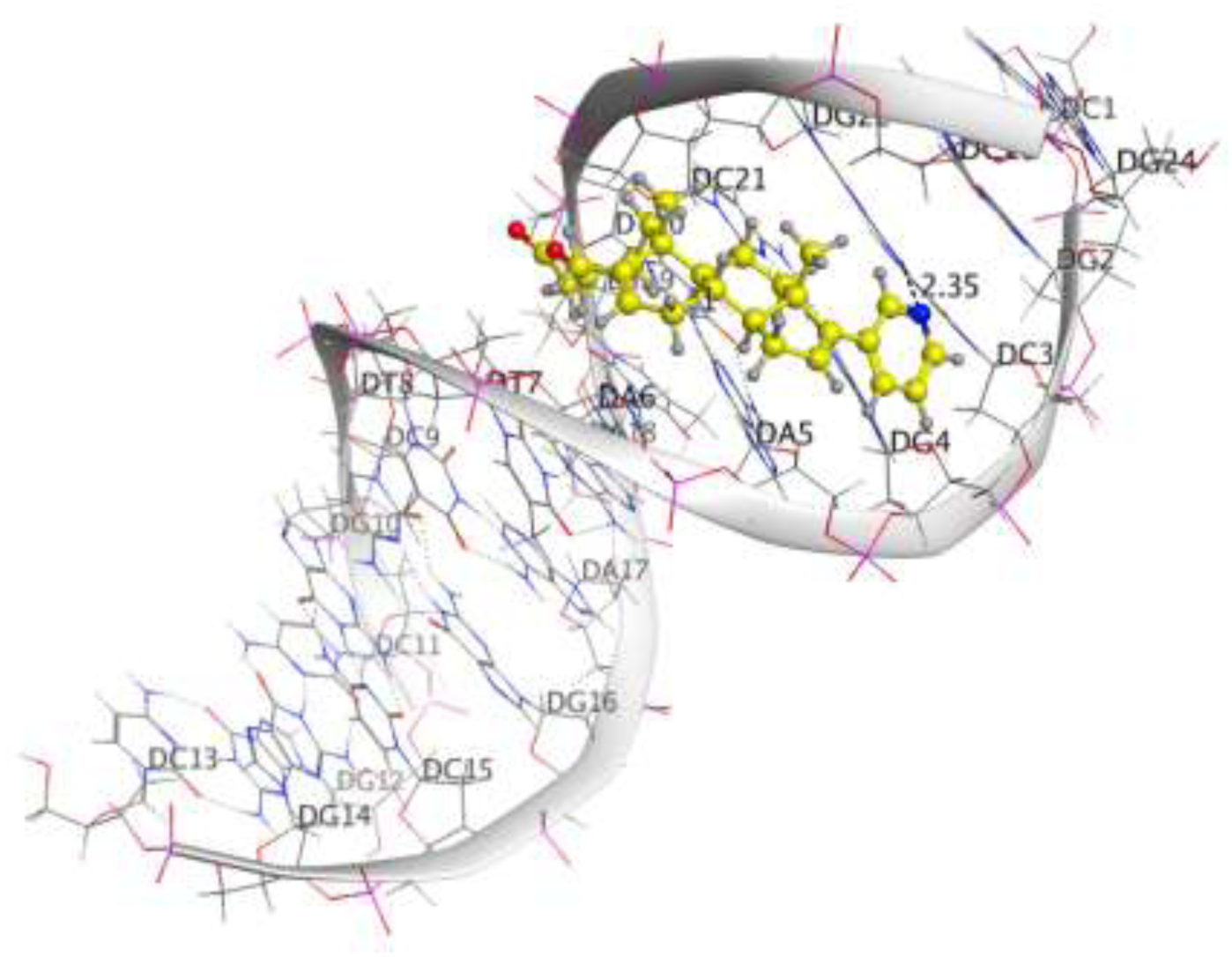
[A] ABR-ctDNA docking conformation with lowest energy. [B] ABR surrounded by DNA bases

## Conclusions

The interaction between ABR and ctDNA indicates the involvement of static quenching. Also the minor groove of ctDNA was found to interact with ABR by spontaneous hydrogen bonding with no other electrostatic interactions being involved in binding mechanism.

## Acknowledgments

The authors would like to extend their sincere appreciation to the Deanship of Scientific Research, King Saud University, for funding the research group No. RG-1438-042.

